# A large-scale assessment of sequence database search tools for homology-based protein function prediction

**DOI:** 10.1101/2023.11.14.567021

**Authors:** Chengxin Zhang, P. Lydia Freddolino

## Abstract

Sequence database searches followed by homology-based function transfer form one of the oldest and most popular approaches for predicting protein functions, such as Gene Ontology (GO) terms. Although sequence search tools are the basis of homology-based protein function prediction, previous studies have scarcely explored how to select the optimal sequence search tools and configure their parameters to achieve the best function prediction. In this paper, we evaluate the effect of using different options from among popular search tools, as well as the impacts of search parameters, on protein function prediction. When predicting GO terms on a large benchmark dataset, we found that BLASTp and MMseqs2 consistently exceed the performance of other tools, including DIAMOND - one of the most popular tools for function prediction - under default search parameters. However, with the correct parameter settings, DIAMOND can perform comparably to BLASTp and MMseqs2 in function prediction. This study emphasizes the critical role of search parameter settings in homology-based function transfer.

## INTRODUCTION

Homology-based function transfer is a classical and widely used approach for predicting protein functions. In this method, the sequence of the query protein is searched through a database of template proteins with function annotations. The biological functions of the query protein are then derived from the annotations of the top sequence search hits. The operation of searching for sequence homologs forms the foundation for many classical protein function predictors [1-5] and remains a critical component for numerous state-of-the-art deep learning-based function prediction algorithms [6-9].

Despite the central role of sequence database search tools in homology-based function prediction, there is no consensus on the best tool and its optimal search parameters for predicting biological functions. While BLASTp [10] was the most commonly used tool by traditional function prediction software [4, 5], more recently developed predictors [7-9] tend to favor DIAMOND [11], a BLASTp alternative optimized for speed. A smaller number of predictors [3, 12] employ PSI-BLAST [10] and MMseqs2 [13]. Interestingly, hidden Markov Model (HMM)-based sequence database search tools such as HHblits [14] and jackhmmer [15], despite their popularity in protein structure prediction [16, 17], are rarely used in function prediction.

Beyond the choice of sequence database search tool, the scoring function used to derive function annotations from a specific set of template proteins also plays a major role in the accuracy of homology-based function prediction. For example, several recent studies [1, 18] consistently indicate that deriving the prediction score from multiple hits tends to produce more accurate predictions than simply deriving the function prediction from the template with the highest sequence identity. However, whether different sequence search tools should employ different scoring functions remains an open question.

To tackle these questions, we performed a direct comparison of popular sequence search tools used for protein structure and function prediction, as well as a set of scoring functions for homology-based function prediction. Working in the context of a large-scale protein Gene Ontology (GO) prediction task, we observed a profound effect of the choices of search tool, tool-specific parameters, and scoring function on the overall performance. Our findings provide a workflow for homology-based function prediction that appears optimal both in terms of accuracy and speed on the datasets used in our comparisons and provide a strong foundation for ongoing efforts in tool development for protein function prediction.

## METHODS

### Protein sequence database search tools

We evaluated seven popular protein sequence database search tools: BLASTp [10], DIAMOND [11], MMseqs2 [13], PSI-BLAST [10], phmmer [15], jackhmmer [15], and HHblits [14]. Among these, BLASTp (version 2.13.0+), DIAMOND (version 2.1.8.162), and MMseqs2 (version 390457d87ed7049d918e46bc8b0571ac4034aae4) are based on sequence-sequence alignment. PSI-BLAST (version 2.13.0+) by default performs sequence-sequence alignment, producing results identical to those from BLASTp. However, rather than using the default setting, we utilized the “-num_iterations 3” option to perform an iterative profile-sequence search with PSI-BLAST. These four programs can utilize a protein sequence database after a straightforward reformatting process to convert the database text into binary formats.

Jackhmmer (version 3.4) and phmmer (version 3.4) are based on HMM-sequence alignment. Both programs can directly search a FASTA format sequence database.

On the other hand, HHblits (version 2.0.15) is based on iterative HMM-HMM alignment and requires a HHblits-format database of HMMs. To construct such a database, we follow our previous protocol [16]. Briefly, all template proteins with GO annotations are grouped into sequence clusters by kClust [19] using a 30% sequence identity cutoff. We then use Clustal Omega [20] to align sequences within each cluster into aligned sequence profiles. These profiles are fed into the hhblitsdb.pl script accompanying the HHblits software to construct the HHblits-format database.

### Scoring functions for function prediction

We implemented 11 different scoring schemes to derive function predictions from a set of template hits identified by sequence database search. In the first scoring function, the score for predicting GO term *q* is:

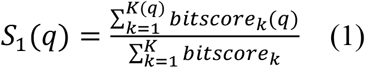

Here, *bitscore*_*k*_ is the bit-score for the *k*-th template; *K* is the total number of templates with GO annotations; *bitscore*_*k*_(*q*) and *K*(*q*) are the respective values for the subset of templates with GO term *q*. This scoring function, where each template is weighted by bit-score, appears to be the most popular scoring function among recently developed machine learning-based function predictors [6, 8, 18, 21]. It was called either BLAST-KNN [18] or DiamondScore [8] by previous studies.

The second scoring function is newly introduced by this study, where the template is weighted by both the bit-score and sequence identity.

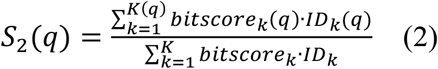

Here, *ID*_*k*_ is the sequence identity for the *k*-th template, calculated by the number of identical residues divided by the maximum between the query sequence length and template sequence length. *ID*_*k*_ (*q*) is the respective value for the *k*-th template with GO term *q*.

The third scoring function was introduced by MetaGO [1], where the template is weighted by *qID*_*k*_, the sequence identity of the *k*-th template normalized by the length of query sequence:

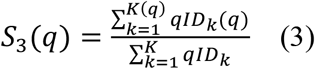

The fourth, fifth and sixth scoring function are defined similarly, except that the sequence identities *tID*., *aID*. and *ID*. are normalized by the length of the *k*-th template, the number of aligned residues, and the maximum between query and template, respectively:

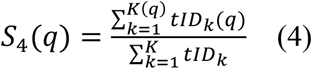

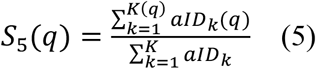

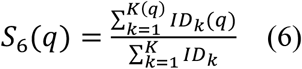

The seventh scoring function is the frequency of templates with GO term *q* among all templates:

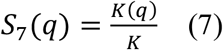

Whereas the previous scoring functions are all calculated from all template hits, the remaining four scoring functions only consider the template with the highest sequence identity among all templates with GO term *q*. In the eighth, ninth, tenth and eleventh scoring functions, the sequence identities are normalized by the query length, template length, number of aligned residues, and the maximum between query length and template length, respectively.

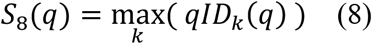

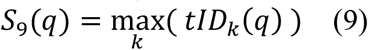

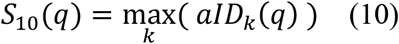

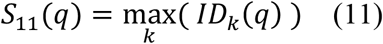

### Evaluation of function prediction performance

The performance of protein function prediction is usually evaluated by the maximum F-measure (Fmax) and/or the maximum of information content-weighted F-measure (wFmax). Fmax was the main evaluation metric for Critical Assessment of Function Annotation (CAFA) challenges round 1, 2 and 3 [22], and is defined as:

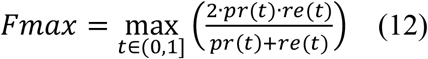

Here, *pr*(*t*) and *re*(*t*) are the precision and recall, respectively, at the prediction score threshold *t*. They are defined as:

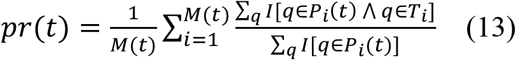

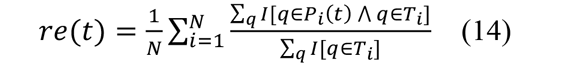

Here, *N* is the total number of proteins in the dataset; *M*(*t*) is the number of proteins with at least one predicted GO term with prediction score ≥ *t. T*_*i*_ is the set of experimentally determined (ground truth) GO terms, including their parent terms, for protein *i. P*_*i*_(*t*) is the set of predicted GO terms for protein *i* with prediction score ≥ *t. I*[] is the standard indicator function (i.e., the Iverson’s bracket). For both *T*_*i*_ and *P*_*i*_(*t*), the root terms of the three GO aspects (GO:0003674 “molecular_function”, GO:0008150 “biological_process”, and GO:0005575 “cellular_component”) are excluded.

The other evaluation metric used here is wFmax, which is the main evaluation metric for the currently-in-progress CAFA5 experiment. It is defined as:

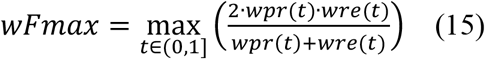

Here, w*pr*(*t*) and w*re*(*t*) are the precision and recall, respectively, weighted by the information contents [23] of GO terms. They are defined as:

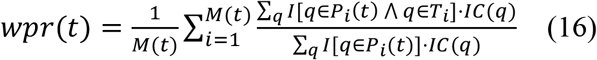

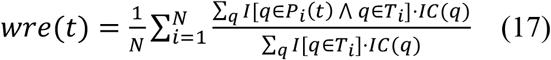

The information content, also known as the information accretion, for GO term *q* is defined as:

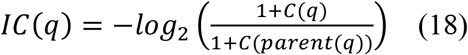

Here, *C*(*q*) is the number of proteins with GO terms q in the whole template database; *C*(*parent(q)*) is the number of database proteins with all parent terms of GO term *q*.

The three aspects of GO – Molecular Function (MF), Biological Process (BP), and Cellular Component (CC) – are evaluated separately.

## RESULTS

### Datasets

We benchmarked different homology-based function prediction schemes on a time-elapsed test set of 4,303 proteins. Each of these proteins has been annotated with at least one new GO term in any one of the three GO aspects in UniProt Gene Ontology Annotation (UniProt-GOA) release 2023-07-12 but has no GO annotation in the same aspect in UniProt-GOA release 2022-11-17. The template sequence database used by the evaluated search tools consists of 134,862 proteins with GO annotations in UniProt-GOA 2022-11-17. In line with the protocols used in recent CAFA challenges, only GO annotations with experimental or high-throughput evidence (evidence codes: EXP, IDA, IPI, IMP, IGI, IEP, HTP, HDA, HMP, HGI, HEP), traceable author statements (evidence code: TAS), or inferences made by curators (evidence code: IC), along with all their parent GO terms, are considered in curating the template database and test set. Proteins whose only leaf MF GO term is GO:0005515 “protein binding” are not included as MF test protein or MF template protein. In total, the test set comprises 1,119 proteins for the MF aspect, 1,609 for the BP aspect, and 2,468 for the CC aspect.

### Scoring functions are critical for function prediction

Our large-scale benchmarks reveal that even when utilizing the same search database tool, varying scoring functions can drastically influence the performance of homology-based protein function prediction (**Figure 1** and **Figure S1**). For instance, when using BLASTp, the top-performing scoring function (*S*_2_) exhibits wFmax values higher by 33.3%, 83.4%, and 15.2% than the worst scoring functions for MF, BP, and CC, respectively.

**Figure 1.**
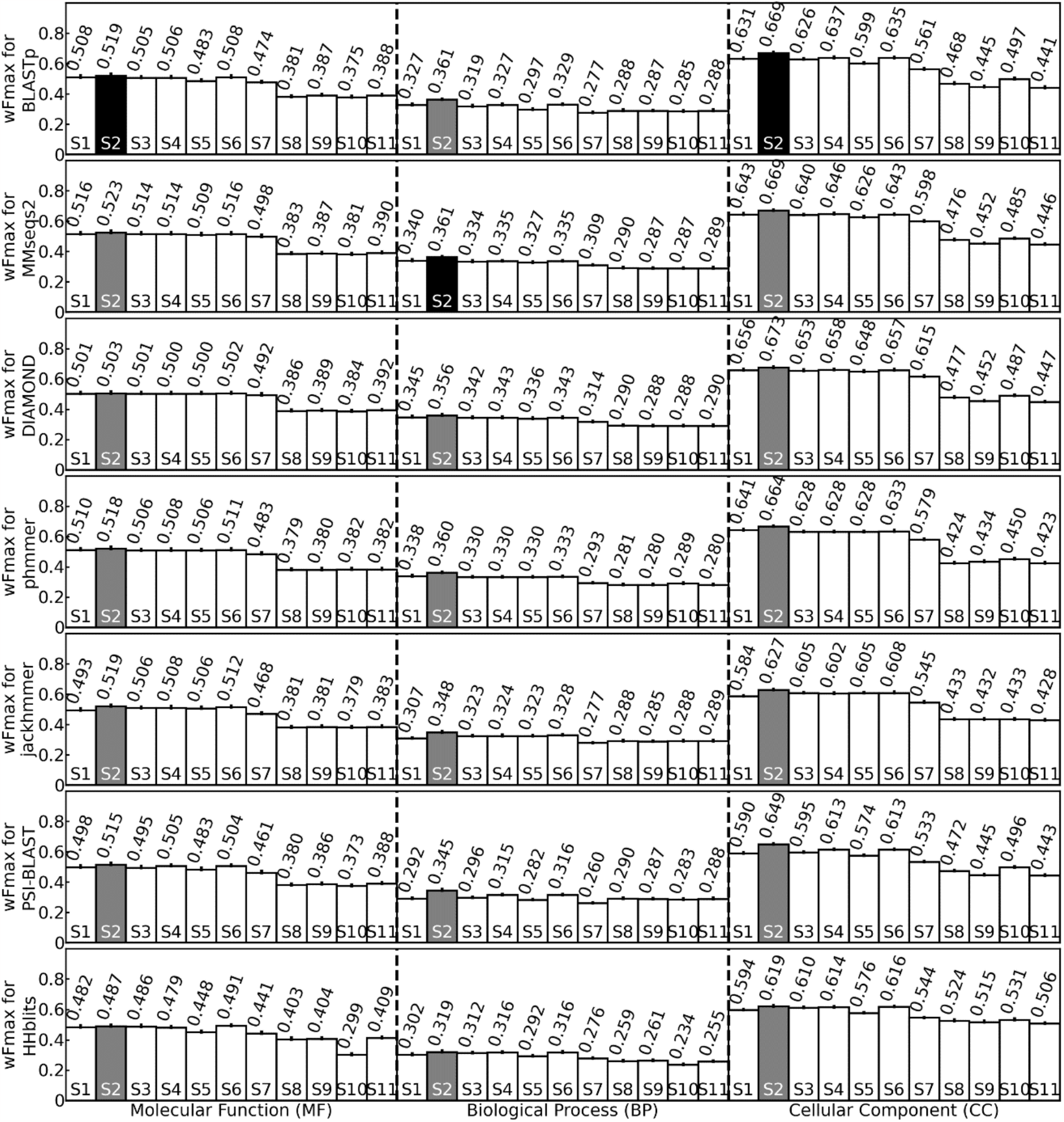
The wFmax values for GO prediction. Each row represents a different database search tool using default parameters, which different bars within each row correspond to the 11 scoring functions. The lengths of the error bars are equal to the standard error of mean (SEM) of weighted F-measure values per protein. Grey bars indicate the highest wFmax value for each method. Black bars indicate the highest wFmax value among all methods for a particular GO aspect.

Independent of sequence database search tools and GO aspects, the scoring function *S*_2_, which weighs all templates by both bit-scores and sequence identities, performs the best (black bars in Figure 1 and Figure S1). This is followed by functions *S*_1_ and *S*_3_ to *S*_6_, which weigh all templates either by bit-scores or sequence identities, but not both. For MF and CC, the worst scoring functions are those that solely consider the template with the highest sequence identity (*S*_8_ to *S*_11_). On the other hand, the scoring function that considers the frequency of a GO term among all templates (*S*_7_) outperforms *S*_8_ to *S*_11_ but still underperforms when compared to *S*_1_ to *S*_6_. For BP, functions *S*_7_ to *S*_11_ exhibit poor performance at similar levels.

### Traditional sequence-sequence alignment outperforms hidden Markov model for function prediction

Our previous study [16] demonstrated that more sensitive HMM-based sequence searches, particularly those by HHblits, markedly enhanced protein structure prediction compared to sequence-sequence alignment. However, the opposite holds true for protein function prediction (**Figure 1**). In fact, HMM-based sequence search tools such as jackhmmer and HHblits generally perform worse than sequence-sequence alignment-based tools like BLASTp and MMseqs2 almost across all scoring functions—with HHblits trailing as the least effective sequence search tool in nearly all scenarios. This discrepancy might be attributed to the optimization of HHblits for structure prediction tasks, including remote structure analog detection [14].

We notice that programs employing iterative database search, e.g., PSI-BLAST and jackhmmer, tend to have worse GO prediction accuracies compared to similar programs using less sensitive non-iterative modes (BLASTp and phmmer, respectively, **Figure 1**). Similarly, for MMseqs2, we observe a minor but consistent decrease in GO prediction accuracies with an increase in the number of iterations (**Figure S2**). This finding contrasts with the case in protein structure prediction tasks [16], where iterative searches typically enhance structure modeling quality, presumably by providing deeper and more informative multiple sequence alignments.

All of these findings unanimously suggest that function prediction necessitates less sensitivity in template detection than structure prediction. This result is not surprising, given previous studies which indicated that even though proteins sharing a sequence identity as low as 30% [24] often exhibit similar structures, a sequence identity of at least 60% [25] must be maintained to ensure similar biological functions. Indeed, more remote homologs may contaminate function predictions with low-confidence information, thus decreasing performance.

Overall, the best GO prediction results are produced by MMseqs2 for MF, by both BLASTp and MMseqs2 for BP, and by DIAMOND for CC. Consequently, the following section will concentrate on these three programs while using the uniformly optimal scoring function (*S*_2_).

### Proper parameter settings improve DIAMOND-based function prediction

While BLASTp, DIAMOND, and MMseqs2 all perform non-iterative sequence-sequence searches, their default settings differ greatly. For instance, the default E-value cutoffs for BLASTp and DIAMOND are 10 and 0.001, respectively, while the default number of top hits are 500 and 25. Moreover, both DIAMOND and MMseqs2 can function under different sensitivity modes. Therefore, we examined how varying search parameters, such as E-value cutoffs, sensitivity modes, and the maximum number of top hits, impact the effectiveness of function prediction.

After assessing various parameter combinations, we discovered that using non-default settings on MMseqs2 only improves function prediction accuracy very slightly (**Figure 2** and **Figure S3**). Conversely, operating BLASTp at a lower E-value cutoff (-evalue 0.1) and reducing the number of hits (-max_target_seqs 100) mildly improves MF, BP, and CC prediction wFmax by 1.3%, 1.9%, and 0.1%, respectively (**Figure 2** and **Figure S4**). DIAMOND benefits the most from parameter tuning. Higher sensitivity settings (--ultra-sensitive, --very-sensitive, or –more-sensitive) generally enhance GO prediction accuracies (**Figure S5** and **Figure S6**). Moreover, a lenient E-value cutoff (--evalue 1) also boosts prediction accuracies (**Figure S7**)

**Figure 2.**
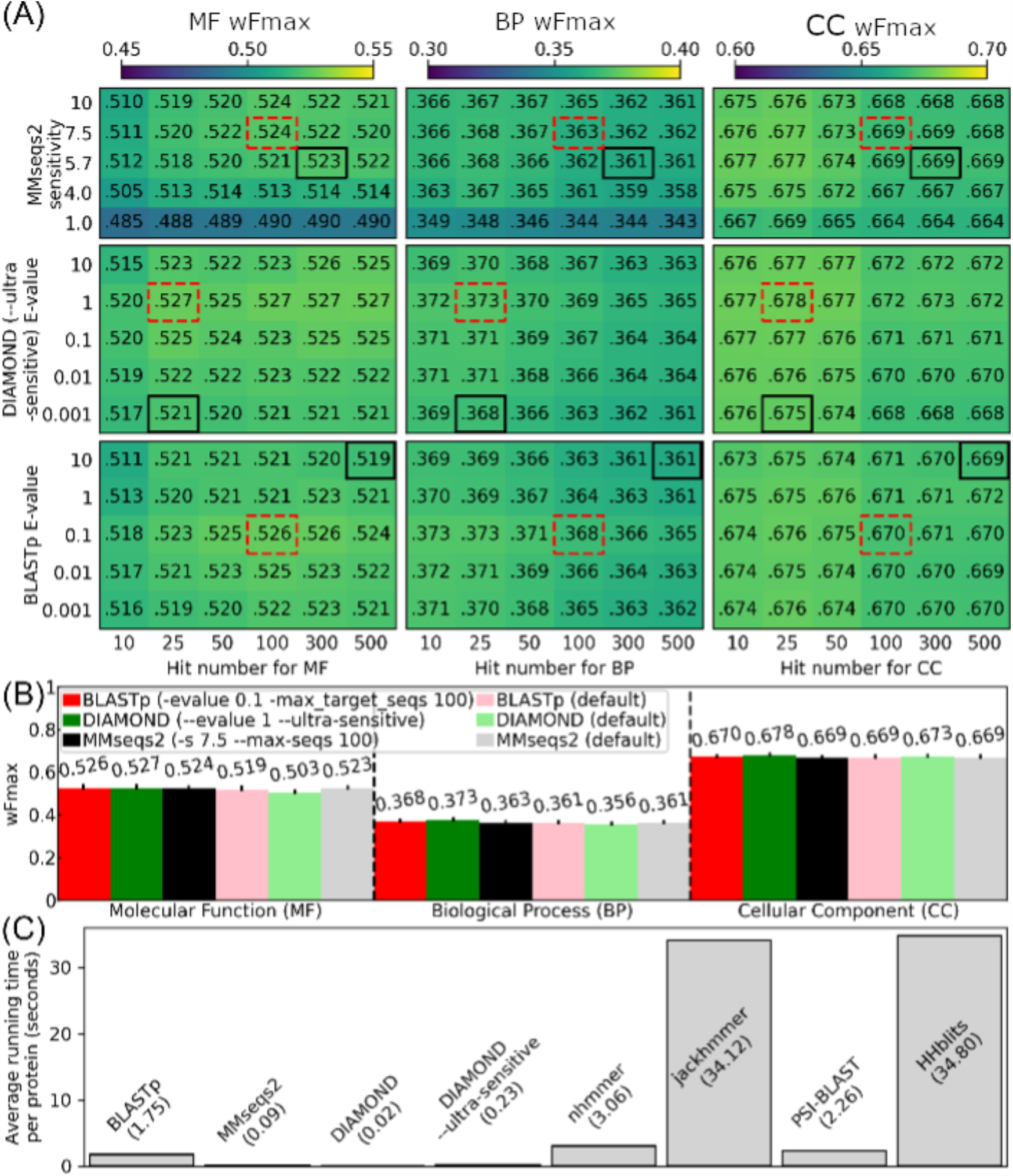
Parameter optimization for GO prediction using scoring function *S*_2_. (**A**) Heatmap for wFmax values using different E-values/sensitivities and maximum hit number. Black solid boxes and red dashed boxes indicate the default and optimized parameters, respectively. (**B**) The wFmax values for the optimal parameters (dark color bars) and default parameters (light color bars) for each tool. The error bar lengths reflect the standard error of mean (SEM) of weighted F-measure values per protein. (**C**) Average running time of different sequence search tools. Since DIAMOND running at the default sensitivity mode and --ultra-sensitive mode have very different speeds, the running times of both are shown.

Despite DIAMOND performing less effectively than MMseqs2 and BLASTp for GO prediction when all programs operate under their default settings, it performs comparably or slightly better than the other two programs when all programs operate with the optimal search parameters (“--evalue 1 --ultra-sensitive” for DIAMOND, “-evalue 0.1 - max_target_seqs 100” for BLASTp, and “-s 7.5 --max-seqs 100” for MMseqs2, **Figure 2**). Specifically, after adjusting search parameters, DIAMOND and BLASTp have comparable MF and BP accuracies, which are superior to that of MMseqs2. With parameter tuning, DIAMOND’s wFmax values for CC shows enhancements of 1.2% over those from BLASTp, which, in turn, outperforms MMseqs2. These adjustments position DIAMOND as an attractive choice for large-scale function prediction, particularly considering its >7 times faster speed compared to BLASTp (**Figure 2C**).

To provide an additional test of our methods on a data set completely separated from our testing to this point, we repeated the same experiment using the CAFA3 dataset (**Figure S8**) and the same set of optimized parameters obtained in **Figure 2B**. The test result on this smaller dataset is largely consistent with the findings above: the GO prediction performance by BLASTp and MMseqs2 are comparable between the default parameters and the optimized parameters. On the other hand, the performance of DIAMOND can be significantly improved by parameter tuning: DIAMOND run under the default setting lags far behind BLASTp and MMseqs2 on all GO aspects but DIAMOND is com-parable to the other program when running at a higher sensitivity mode.

### Case study: function prediction of *Drosophila melanogaster* TMTC4

To delve further into the impact of different sequence search tools on function prediction, we use the TMTC4 protein from the fruit fly (UniProt accession: Q9VF81) as a case study. This protein is a dolichyl-phosphate-mannose-protein mannosyltransferase (GO:0004169) [26] (**Figure 3A**). While none of the programs can predict this highly specific MF GO term (**Figure 3B**), all of them can predict some parent terms of this GO term. DIAMOND and MMseqs2 provide the most specific parent term, GO:0000030 “mannosyltransferase activity” (Figure 3C). Next in line is BLASTp (**Figure 3D**), followed by HHblits and phmmer (**Figure 3E**) predicting its parent term, GO:0016758 “hexosyltransferase activity”. The least specific predictions come from PSI-BLAST and jackhmmer (**Figure 3F**), which merely predict an even less specific parent term, GO:0016757 “glycosyltransferase activity”.

**Figure 3.**
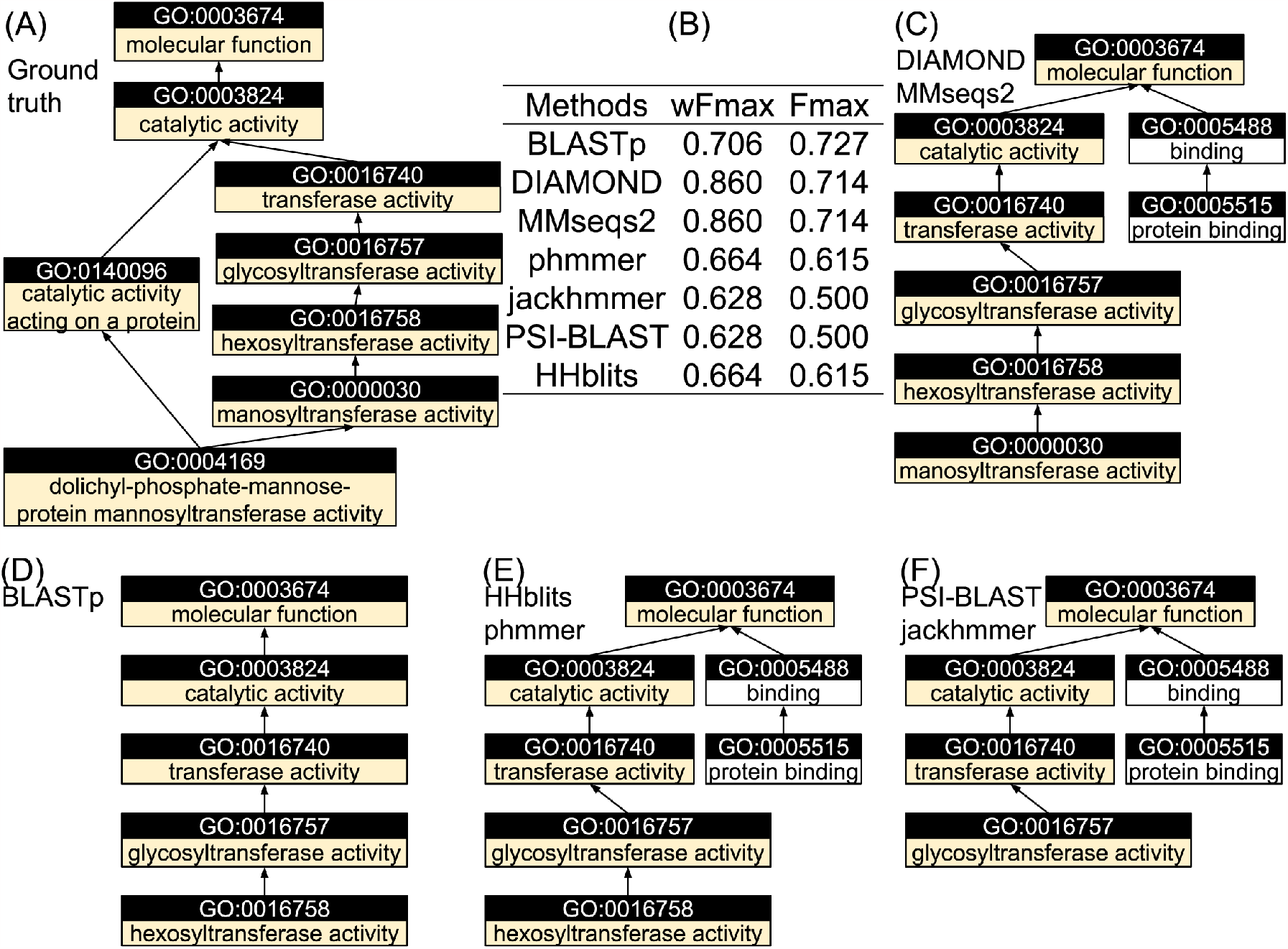
MF GO annotation for TMTC4 (UniProt accession: Q9VF81). (**A**) Ground truth annotation. (**B**) Weighted F-measure and F-measure of MF prediction by different tools. (**C-F**) GO predictions by (**C**) DIAMOND and MMseqs2, (**D**) BLASTp, (**E**) HHblits and phmmer, and (**F**) PSI-BLAST and jackhmmer. Only GO terms predicted above the cutoff corresponding to the maximum weighted F-measure are shown.

The inferior performance of the iterative searching tools (PSI-BLAST and jackhmmer) relative to their respective non-iterative counterparts (BLASTp and phmmer) primarily results from incorporating functionally less relevant templates during the iterations. For instance, in jackhmmer’s first iteration, 107 out of 344 (31.1%) hits are annotated with the correct parent term, GO:0016758. This ratio decreases to 89 out of 1273 (7.0%) and 75 out of 1767 (4.2%) hits in the second and final iterations, respectively. Consequently, the prediction score of GO:0016758 plunges from 0.777 in the first iteration to 0.460 in the final iteration. This phenomenon, known as “profile drift” [27], implies that an increased number of iterations progressively change the composition of the query sequence profile, leading to the inclusion of very distant sequence relatives in the search results.

## CONCLUSIONS

In this study, we evaluated the usefulness of several commonly used sequence search tools for protein function prediction. We discovered that, despite DIAMOND being less effective than BLASTp and MMseqs2 for GO prediction under default parameter settings, it can surpass both programs in terms of GO prediction accuracies merely by adjusting the sensitivity and E-value cutoff settings. These three methods demonstrate higher GO prediction accuracies compared to more sensitive sequence search protocols, including PSI-BLAST, HHblits, jackhmmer, and phmmer, all of which are based on HMMs, iterative search, or both. Alongside evaluating different search tools, this study also reaffirms previous findings that GO predictions stemming from multiple templates are more accurate than those derived from the template with the highest sequence identity. We also identified a new scoring function (*S*_2_) that consistently outshines the extensively employed DiamondScore (*S*_1_) scoring function featured in many function prediction programs [6-8, 18, 21].

In this study, we did not evaluate whether the combination of multiple sequence search tools could yield further improved function prediction accuracies; this remains a valuable topic for future investigation.

## ACKNOWLEDGEMENTS

We thank Quancheng Liu and Dr. Xiaoqiong Wei for insightful discussions. This work used the Advanced Cyberinfrastructure Coordination Ecosystem: Services & Support (ACCESS) program, which is supported by the National Science Foundation (2138259, 2138286, 2138307, 2137603, and 2138296). This work has been supported by the National Institutes of Health (AI134678 to P.L.F.).

## REFERENCES

1. Zhang, C.X., et al., MetaGO: Predicting Gene Ontology of Non-homologous Proteins Through Low-Resolution Protein Structure Prediction and Protein Protein Network Mapping. Journal of Molecular Biology, 2018. 430(15): p. 2256–2265.

2. Zhang, C., P.L. Freddolino, and Y. Zhang, COFACTOR: improved protein function prediction by combining structure, sequence and protein-protein interaction information. Nucleic Acids Res, 2017. 45(W1): p. W291–W299.

3. Gong, Q., W. Ning, and W. Tian, GoFDR: A sequence alignment based method for predicting protein functions. Methods, 2016. 93: p. 3–14.

4. Conesa, A. and S. Gotz, Blast2GO: A comprehensive suite for functional analysis in plant genomics. Int J Plant Genomics, 2008. 2008: p. 619832.

5. Martin, D.M., M. Berriman, and G.J. Barton, GOtcha: a new method for prediction of protein function assessed by the annotation of seven genomes. BMC Bioinformatics, 2004. 5: p. 178.

6. Zhu, Y.H., et al., Integrating unsupervised language model with triplet neural networks for protein gene ontology prediction. PLoS Comput Biol, 2022. 18(12): p. e1010793.

7. Cao, Y. and Y. Shen, TALE: Transformer-based protein function Annotation with joint sequence-Label Embedding. Bioinformatics, 2021. 37(18): p. 2825–2833.

8. Kulmanov, M. and R. Hoehndorf, DeepGOPlus: improved protein function prediction from sequence. Bioinformatics, 2021. 37(8): p. 1187.

9. Yuan, Q., et al., Fast and accurate protein function prediction from sequence through pretrained language model and homology-based label diffusion. Briefings in Bioinformatics, 2023. 24(3).

10. Altschul, S.F., et al., Gapped BLAST and PSI-BLAST: a new generation of protein database search programs. Nucleic Acids Res, 1997. 25(17): p. 3389–402.

11. Buchfink, B., K. Reuter, and H.-G. Drost, Sensitive protein alignments at tree-of-life scale using DIAMOND. Nature methods, 2021. 18(4): p. 366–368.

12. Mahlich, Y., et al., HFSP: high speed homology-driven function annotation of proteins. Bioinformatics, 2018. 34(13): p. i304–i312.

13. Steinegger, M. and J. Soding, MMseqs2 enables sensitive protein sequence searching for the analysis of massive data sets. Nat Biotechnol, 2017. 35(11): p. 1026–1028.

14. Remmert, M., et al., HHblits: lightning-fast iterative protein sequence searching by HMM-HMM alignment. Nature Methods, 2012. 9(2): p. 173–175.

15. Eddy, S.R., Profile hidden Markov models. Bioinformatics, 1998. 14(9): p. 755–63.

16. Zhang, C.X., et al., DeepMSA: constructing deep multiple sequence alignment to improve contact prediction and fold-recognition for distant-homology proteins. Bioinformatics, 2020. 36(7): p. 2105–2112.

17. Jumper, J., et al., Highly accurate protein structure prediction with AlphaFold. Nature, 2021. 596(7873): p. 583–589.

18. You, R., et al., GOLabeler: improving sequence-based large-scale protein function prediction by learning to rank. Bioinformatics, 2018. 34(14): p. 2465–2473.

19. Hauser, M., C.E. Mayer, and J. Soding, kClust: fast and sensitive clustering of large protein sequence databases. BMC Bioinformatics, 2013. 14: p. 248.

20. Sievers, F., et al., Fast, scalable generation of high-quality protein multiple sequence alignments using Clustal Omega. Molecular Systems Biology, 2011. 7.

21. You, R., et al., NetGO: improving large-scale protein function prediction with massive network information. Nucleic Acids Res, 2019. 47(W1): p. W379–W387.

22. Zhou, N.H. and et al, The CAFA challenge reports improved protein function prediction and new functional annotations for hundreds of genes through experimental screens. Genome Biology, 2019. 20(1).

23. Clark, W.T. and P. Radivojac, Information-theoretic evaluation of predicted ontological annotations. Bioinformatics, 2013. 29(13): p. i53–61.

24. Xiang, Z., Advances in homology protein structure modeling. Curr Protein Pept Sci, 2006. 7(3): p. 217–27.

25. Tian, W. and J. Skolnick, How well is enzyme function conserved as a function of pairwise sequence identity? Journal of molecular biology, 2003. 333(4): p. 863–882.

26. Eisenhaber, B., et al., Conserved sequence motifs in human TMTC1, TMTC2, TMTC3, and TMTC4, new O-mannosyltransferases from the GT-C/PMT clan, are rationalized as ligand binding sites. Biology Direct, 2021. 16(1): p. 1–18.

27. Kandathil, S.M., J.G. Greener, and D.T. Jones, Prediction of interresidue contacts with DeepMetaPSICOV in CASP13. Proteins, 2019. 87(12): p. 1092–1099.

