## Supplemental Material for "A large-scale assessment of sequence database search tools for homology-based protein function prediction"

#### Supporting Information

#### Table of Content

**Supporting Figures**

**Figure S1.** The Fmax values for GO prediction by 7 database search tools and 11 scoring functions.

**Figure S2.** The wFmax values for GO prediction by MMseqs2 using different number of iterations and different number of hits.

**Figure S3.** The wFmax and Fmax values for GO prediction by MMseqs2 using different sensitivities and hit numbers.

**Figure S4.** The wFmax and Fmax values for GO prediction by BLASTp using different E-value cutoffs and hit numbers.

**Figure S5.** The wFmax values for GO prediction by DIAMOND using 11 scoring functions and different sensitivity levels.

**Figure S6.** The Fmax values for GO prediction by DIAMOND using 11 scoring functions and different sensitivity levels.

**Figure S7.** The wFmax and Fmax values for GO prediction by DIAMOND using different E-value cutoffs and hit numbers.

**Figure S8.** The wFmax values for GO prediction on the CAFA3 set using the optimized parameters and default parameters for each tool.

### Supporting Figures

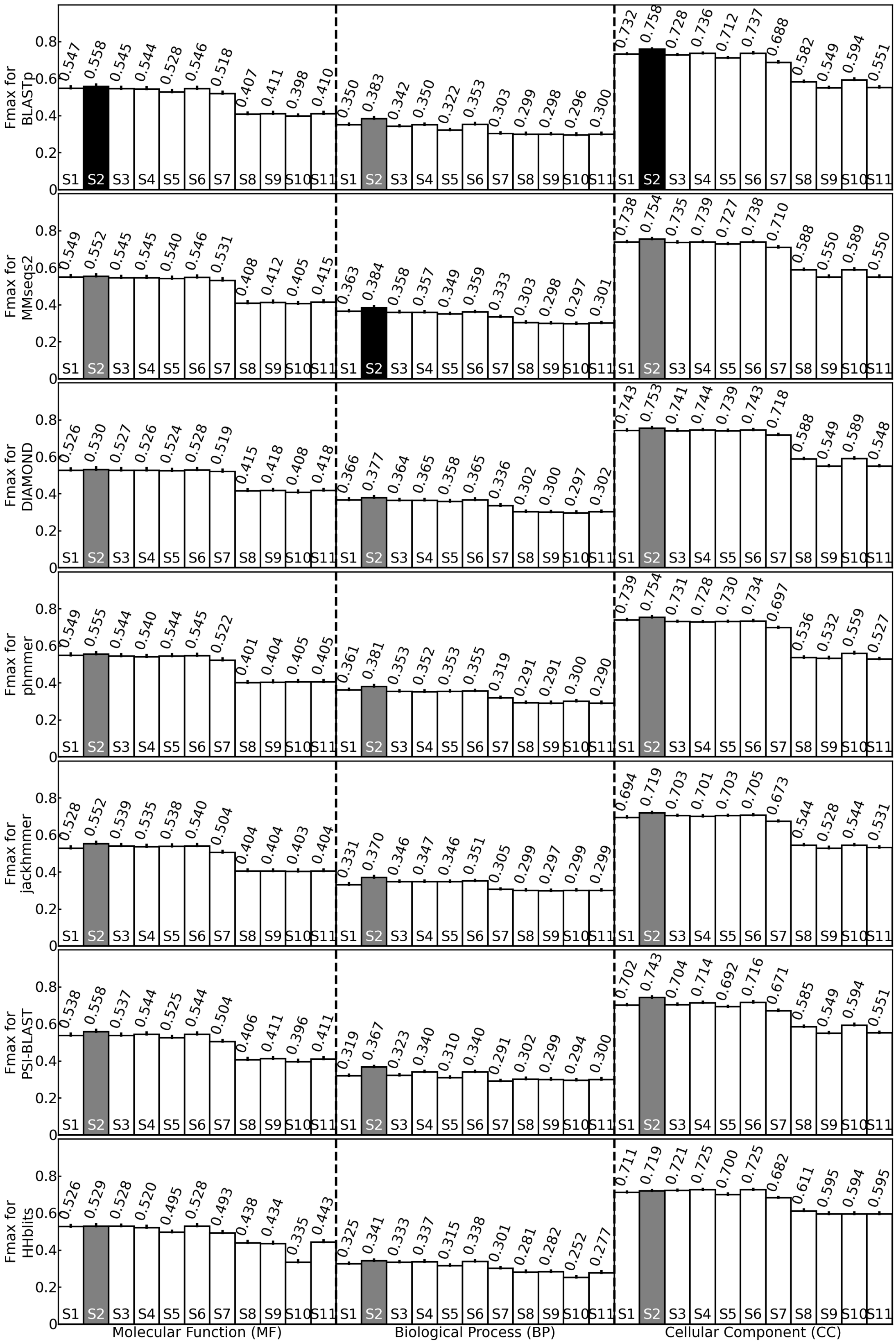


**Figure S1.** The Fmax values for GO prediction by 7 database search tools (different rows) and 11 scoring functions (different bars in a row). Error bars indicate standard error of mean (SEM) of the per protein Fmax values. Grey bars indicate the highest Fmax value for each method. Black bars indicate the highest Fmax value among all methods.

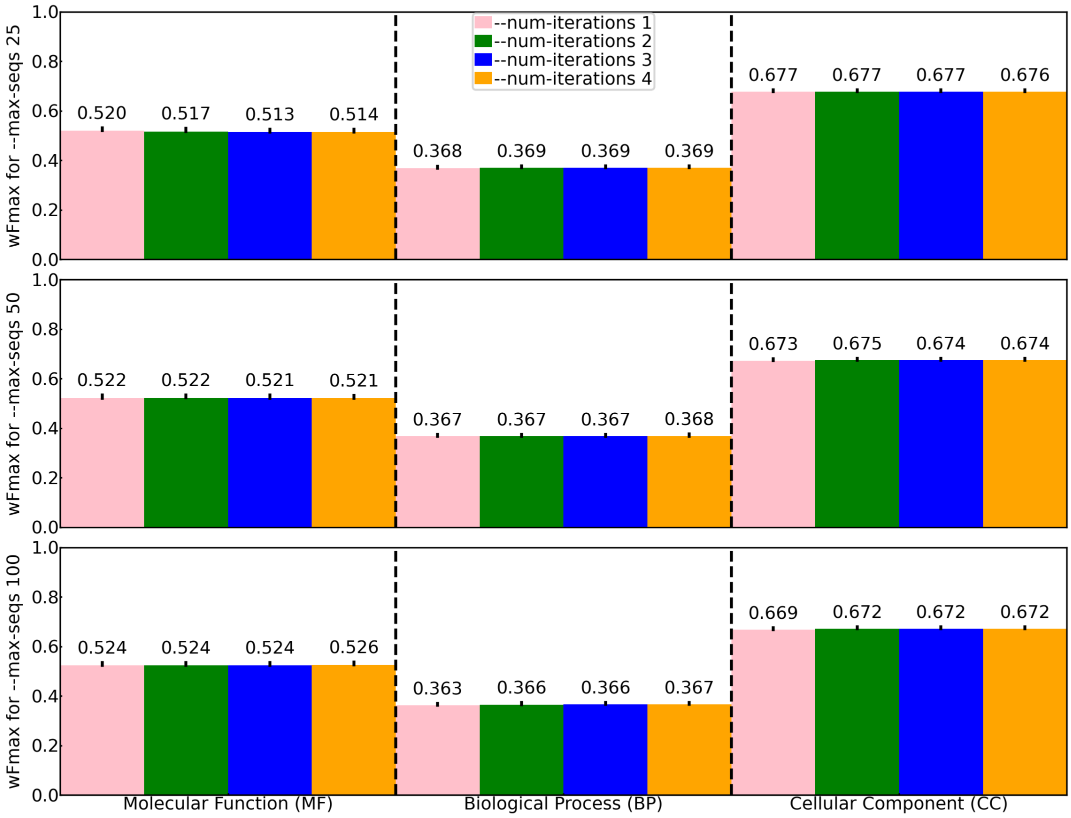


**Figure S2.** The wFmax values for GO prediction by MMseqs2 using different number of iterations (different bars) and different numbers of hits (different rows). The lengths of the error bars equal to the standard error of mean (SEM) of the per protein wFmax values.

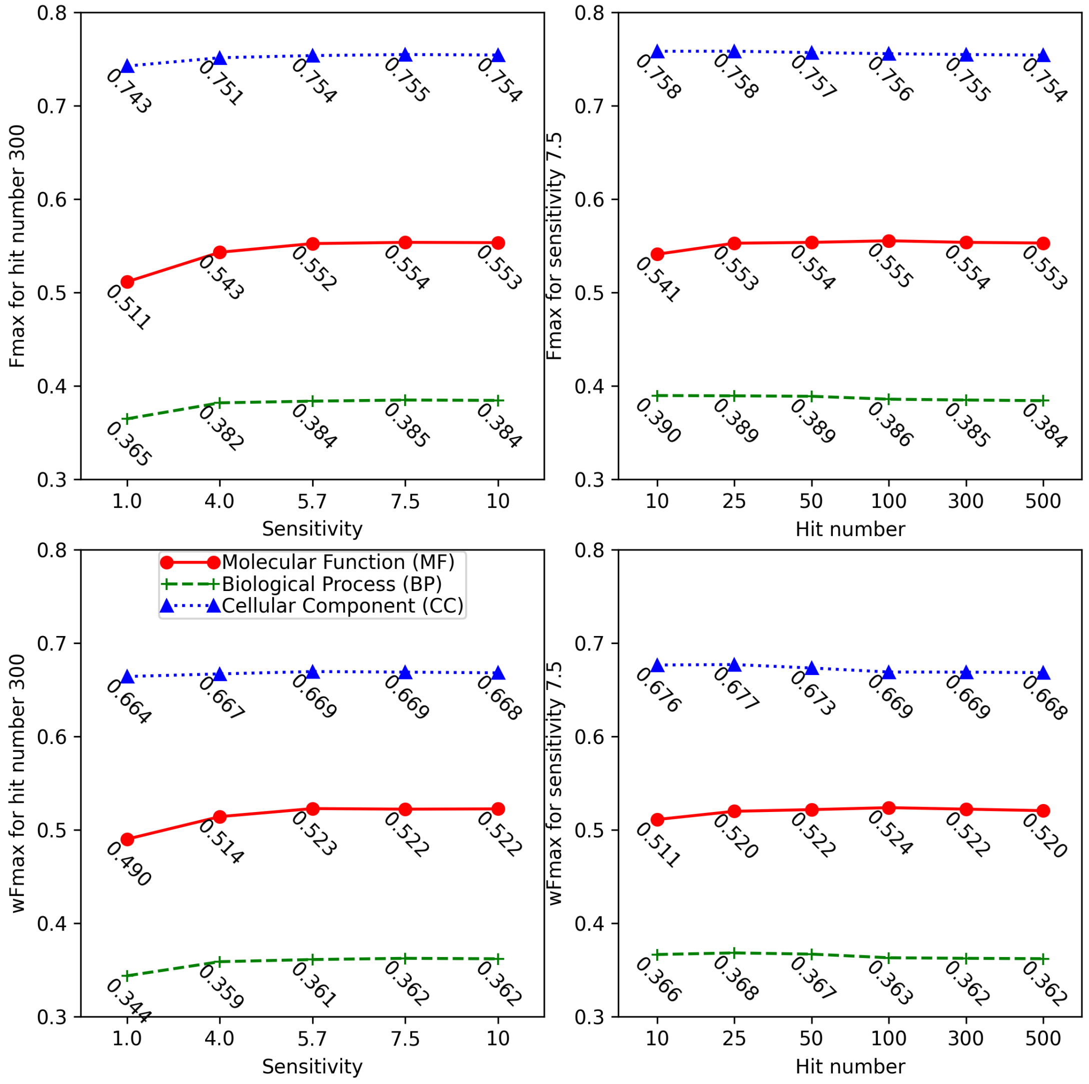


**Figure S3.** The wFmax and Fmax values for GO prediction by MMseqs2 using different sensitivities (-s) and hit numbers (--max-seqs). Here, hit number 300 and sensitivity 5.7 are the default values used by MMseqs2.

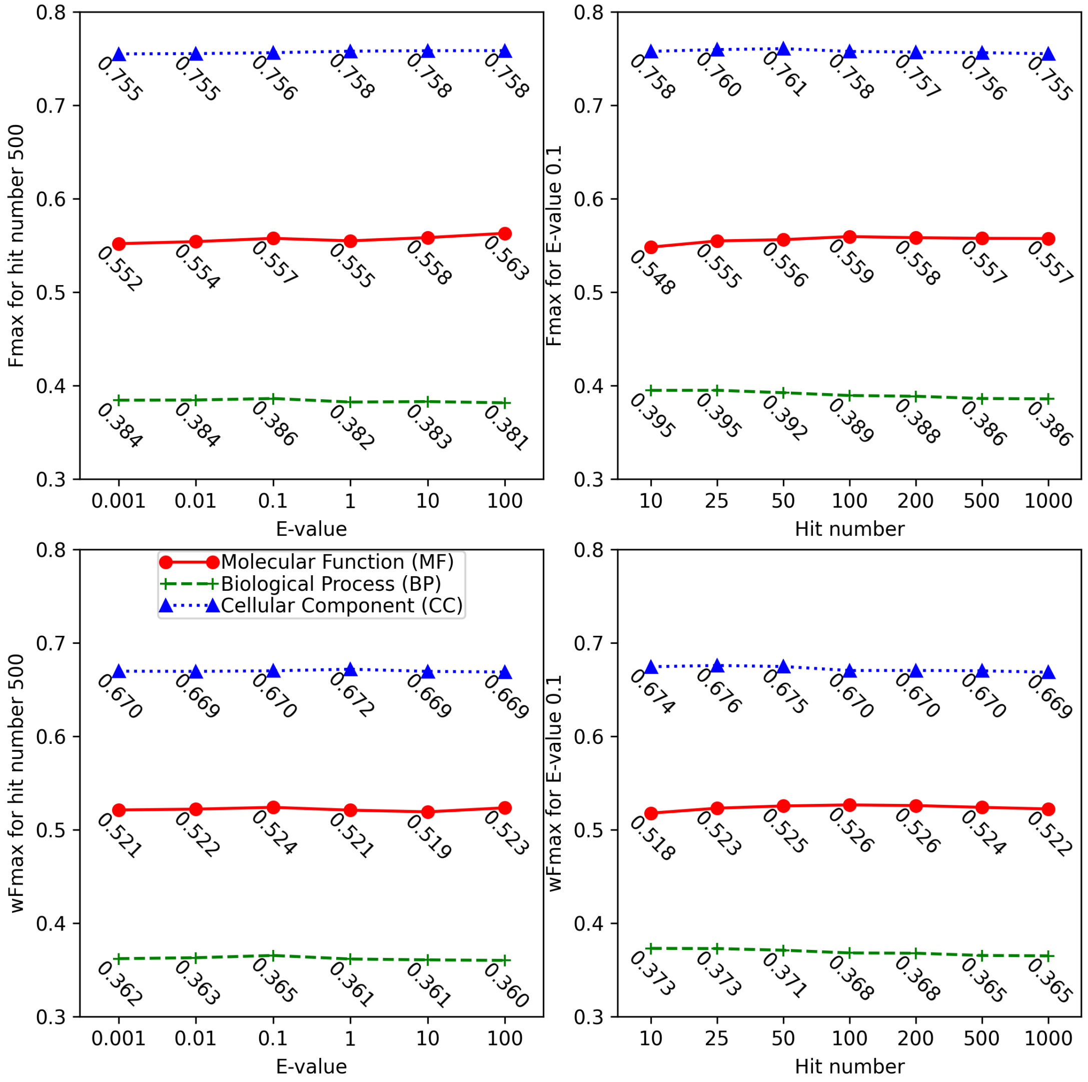


**Figure S4.** The wFmax and Fmax values for GO prediction by BLASTp using different E-value cutoffs (-evalue) and hit numbers (-max_target_seqs). Here, hit number 500 is the default value used by BLASTp. E-value 0.1 is the optimal cutoff to achieve the best GO prediction accuracy.

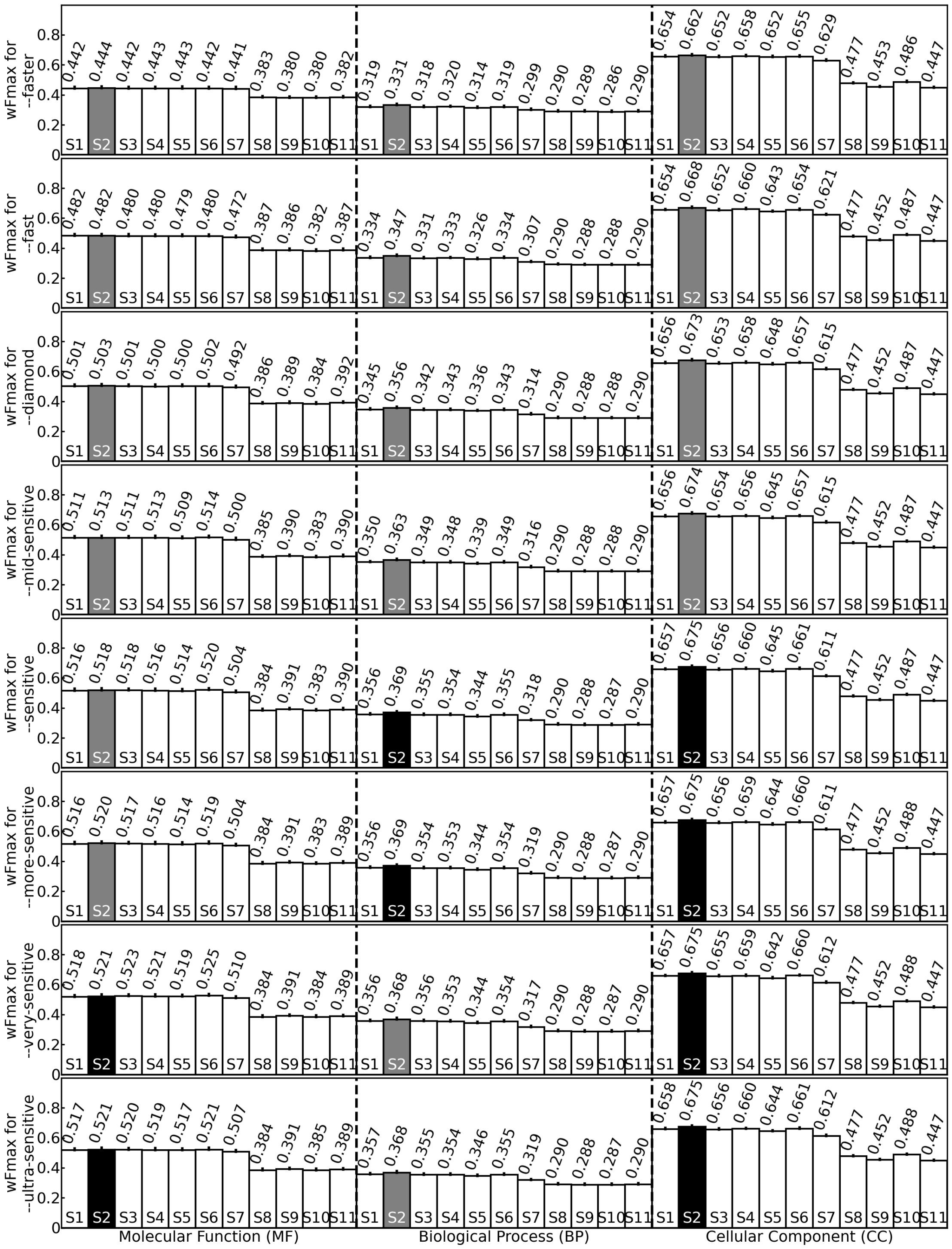


**Figure S5.** The wFmax values for GO prediction by DIAMOND using 11 scoring functions (different bars in a row) and different sensitivity levels (different rows). The lengths of the error bars equal to the standard error of mean (SEM) of the per protein wFmax values. Grey bars indicate the highest wFmax value for each DIAMOND sensitivity mode. Black bars indicate the highest wFmax value for that GO aspect among all DIAMOND sensitivity modes.

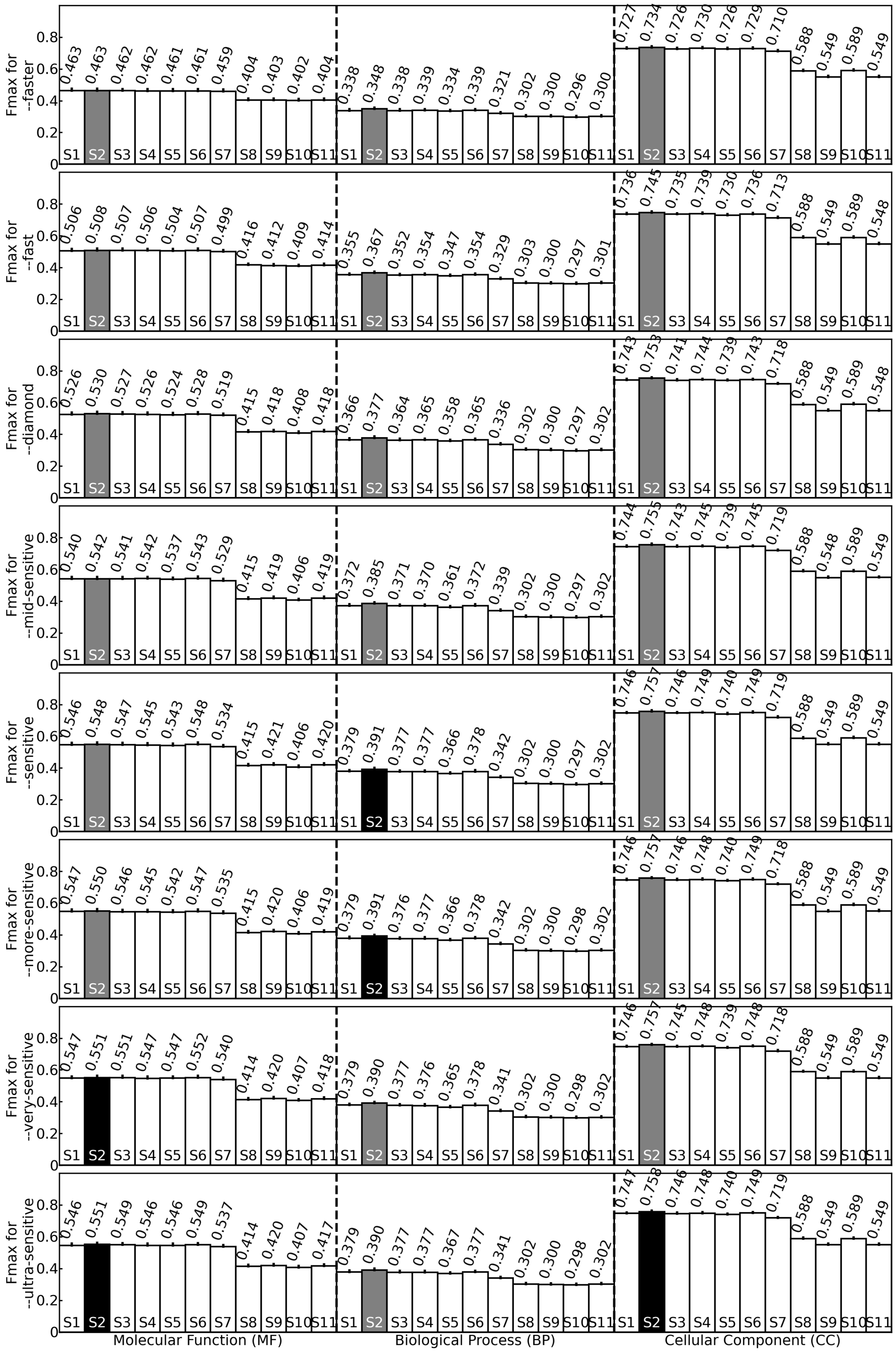


**Figure S6.** The Fmax values for GO prediction by DIAMOND using 11 scoring functions (different bars in a row) and different sensitivity levels (different rows). The lengths of the error bars equal to the standard error of mean (SEM) of the per protein Fmax values. Grey bars indicate the highest Fmax value for each DIAMOND sensitivity mode. Black bars indicate the highest Fmax value for that GO aspect among all DIAMOND sensitivity modes.

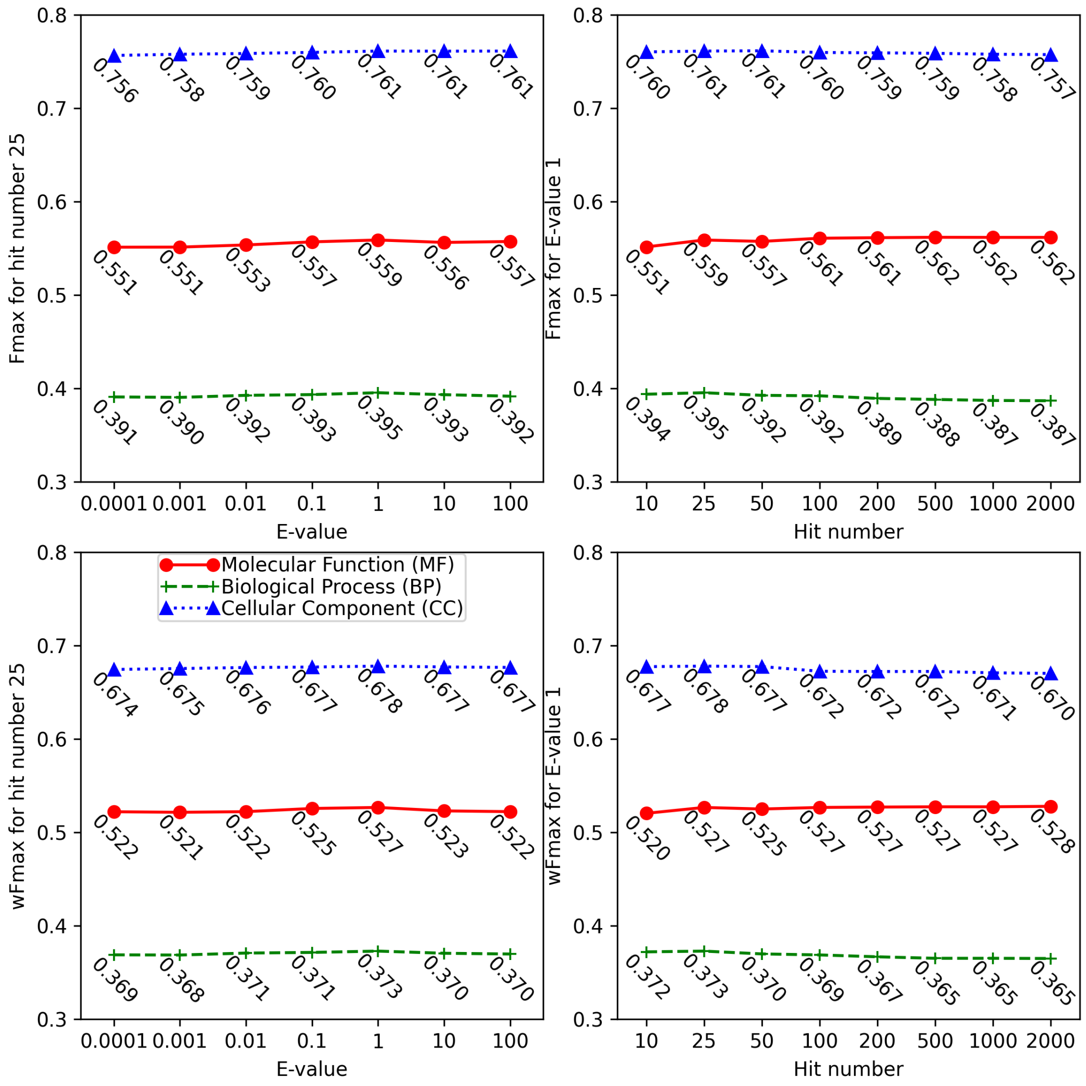


**Figure S7.** The wFmax and Fmax values for GO prediction by DIAMOND using different E-value cutoffs (--evalue) and hit numbers (--max-target-seqs). Here, hit number 25 is the default value used by DIAMOND. E-value 1 is the optimal cutoff to achieve the best GO prediction accuracy.

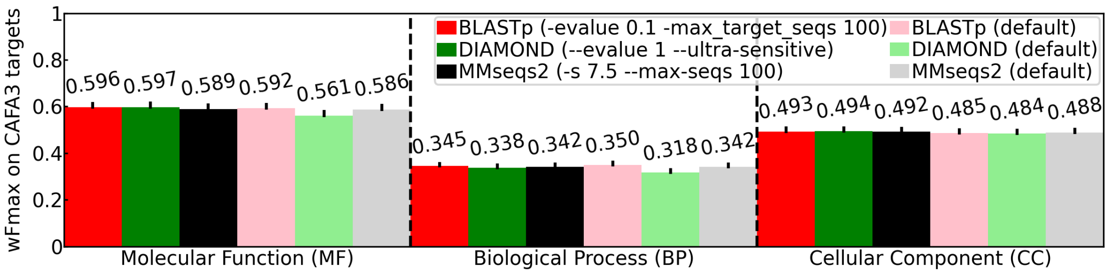


**Figure S8.** The wFmax values for GO prediction on the CAFA3 set using our optimized parameters (optimized on the main benchmark dataset, dark color bars) and default parameters (light color bars) for each tool. The error bar lengths reflect the standard error of mean (SEM) of weighted F-measure values per protein. The CAFA3 dataset consists of 66841 training proteins with GO annotations before September 2016, and 1222 testing proteins with new GO terms annotated on or before June 2017 but after January 2017.
